# The ambivalent effect of spatial structure on the spread of cooperative anti-CRISPR phages

**DOI:** 10.1101/2025.08.06.668856

**Authors:** Anne Chevallereau, Ryuichi Kumata, Olivier Fradet, Sébastien Lion, Edze R. Westra, Akira Sasaki, Sylvain Gandon

## Abstract

Some phages have evolved the ability to cooperate to evade the immunity triggered by their bacterial host. A first exposure to the phage may weaken the host defences and allow later infections to be successful. Because this cooperation requires sequential infections, the phage can invade the host population only if its initial density is sufficiently high in a well-mixed environment. However, most natural bacterial populations are spatially structured. Could spatial structure create more favourable conditions for viral spread? Here we study the effect of spatial structure on the dynamics of cooperative anti-CRISPR (Acr) phages spreading in a population of CRISPR-Cas resistant *Pseudomonas aeruginosa* bacteria. We show experimentally that spatial structure does not always promote the spread of Acr-phages. In particular, the effect of spatial structure is modulated by the efficacy of the bacterial host’s CRISPR-Cas resistance and by the efficacy of the phage Acr protein. These results are discussed in the light of a mathematical model we developed to describe the spread of the phage. The model allows us to understand the ambivalent effects of spatial structure via its effects on the reproduction and on the persistence of the phage. More generally, we find that spatial structure can have opposite effects on the epidemiological dynamics of the phage, depending on the properties of the Acr protein encoded by the phage. This joint experimental and theoretical work yields a deeper understanding of the spatial dynamics of cooperative strategies in phages.

## Introduction

Bacteriophages (phages), the virus that infect bacteria, play a key role in shaping microbial community composition and function [1]. To protect themselves against phages, bacteria have evolved defence strategies and hundreds of novel mechanisms have been uncovered in the past few years [2]. Reciprocally, phages evolved escape strategies to bypass these defence systems by acquiring point (epi)genetic mutations or by encoding inhibitory proteins [3]. Mirroring the accelerated discovery of defence systems, dozens of phage-encoded anti-defence proteins have been identified recently such as anti-CRISPR [4], anti-CBASS [5], anti-Pycsar [5], anti-Thoeris [6,7], etc. While the molecular mechanisms and the biochemistry of these systems are intensely studied, there is an important gap in our understanding of how ecological factors and population dynamics can influence the interactions between bacterial defences and phage counter-defences [8]. Yet, these effects can have major implications on the spread of the phage population. For example, *Pseudomonas aeruginosa* phages encoding anti-CRISPR (Acr) can only amplify if their initial concentration is above a density threshold because they have to cooperate to bypass the CRISPR-Cas defence system of their host [9,10]. This cooperation occurs because Acr proteins only partially protect phages from CRISPR-Cas activity and it takes several rounds of sequential Acr-phage infections to lyse a CRISPR-resistant bacterium. Mechanistically it has been proposed that the first infection by an Acr-phage, although unsuccessful, allows for the delivery of Acr proteins within the host cell which enters a transitory immunosuppressed state. If the Acr-phage density is high, the immunosuppressed host can be reinfected with a second Acr-phage which can now complete its infection cycle and yield progeny. If Acr-phage density is low (i.e. short-term reinfection is unlikely) the immunosuppressed host will revert to a fully resistant state. In other words, this cooperative strategy of the phage within the infected host cell generates a density-dependent effect at the scale of the population (i.e. an Allee effect).

This phage-phage cooperation has previously been explored via experiments carried out in well-mixed conditions. But the dynamics of phage cooperation in natural microbial populations, which are generally structured in space, has received less attention. When reproduction and dispersal are localized, cooperative individuals are more likely to interact with one another than would be expected by chance alone in a well-mixed environment. This clustering of cooperators increases the likelihood that cooperative behaviours will be reciprocated, thereby promoting the persistence and spread of cooperation. As such, spatial structure plays a fundamental role in the evolution and stability of cooperative strategies [11–15]. One may thus expect the spread of Acr-phages to increase in spatially structured environments.

We tested this prediction using first an experimental approach to track how increasing levels of spatial structure of the environment affects the spread of Acr-phages. These experiments yielded unexpected results in which spatial structure can either promote or hinder the spread of the phage population, depending on the strength of the CRISPR-Cas resistance of the bacteria and the efficacy of the Acr mechanism of the phage. Second, to make sense of these results, we extended previous theoretical analyses of the dynamics of Acr-phages to investigate the spread of the phage in a spatially structured environment. This model allowed us to make several new predictions on the effect of key properties of bacterial resistance and Acr efficacy on the spread of Acr-phages. Finally, we went back to the experimental results and checked the validity of these theoretical predictions, confirming the ability of this new model to generate insights and in particular to explain the ambivalent effect of spatial structure on the spread of Acr-phages.

## Results

### Spatial structure favours the growth of DMS3mvir-*acrIF1*

As a protection against phages, bacteria with CRISPR-Cas systems can incorporate short sequences perfectly matching the phage genome, called spacers, into a CRISPR array. The transcript, corresponding to a single spacer, associates with Cas proteins to form a complex that can recognize and cleave matching phage nucleic acid. The level of resistance of the strain against a phage increases with the number of targeting spacers. A strain equipped with spacers, and hence resistant to a phage, is referred to as Bacteriophage Insensitive Mutant (BIM). Phages equipped with Acr can bypass recognition or cleavage and therefore productively infect BIM bacteria. To experimentally explore the effect of spatial structure on the interactions between bacteria with CRISPR-Cas and phages with Acr, we used the model system of *P. aeruginosa* strain PA14 carrying a type I-F CRISPR-Cas system, or a mutant strain with a functionally impaired CRISPR-Cas system (referred to as CRISPR-KO). This strain can be infected with phage DMS3mvir encoding the anti-CRISPR protein AcrIE3 or isogenic mutants encoding AcrIF1 or AcrIF4. Note that AcrIE3 does not inhibit the activity of type I-F CRISPR system and therefore we refer to wildtype DMS3mvir as Acr(-).

We previously showed that upon a primary infection of BIM bacteria, phages equipped with AcrIF1 generate more productive infections than phages encoding AcrIF4, and we refer to these phages as strong and weak Acr-phages, respectively [16]. To manipulate the level of spatial structure in our experiments, we added different concentrations of agar from 0 (liquid) to 1% (solid) to M9 minimal medium. We first validated that we could reproducibly extract and count phages in these settings (**Figure S1**).

To test how increasing the level of spatial structure affects the epidemiological dynamics of Acr-phages, phage-sensitive (CRISPR-KO) or phage-resistant (BIM) strains were infected with phages in 3 media with varying levels of spatial structure (0, 0,5 or 1% agar). Three other parameters were also manipulated: phage initial multiplicities of infection (from 0,001 to 10), levels of resistance (BIM equipped with 3 or 5 targeting spacers, denoted BIM-3sp. and BIM-5sp., respectively) and level of Acr protection (using phage DMS3mvir lacking a functional Acr or encoding weak AcrIF4 or strong AcrIF1)). One day post infection (dpi), phages were extracted and their amplification (ratio of final to initial concentrations) was measured (**Figure 1**). As expected, when CRISPR-Cas is impaired (CRISPR-KO strain) phages amplify regardless of their initial density (**Figure 1A, D, G**, all lines are above the dotted horizontal line). Interestingly, the magnitude of this amplification is similar in all spatial treatments, showing that spatial structure has no effect on Acr-phage epidemiology in absence of CRISPR-resistance. The decrease in amplification with increasing inoculum reflects the fact that phages reach their carrying capacity (approximately 10^9^ pfu/ml) within one day after infection.

**Figure 1.**
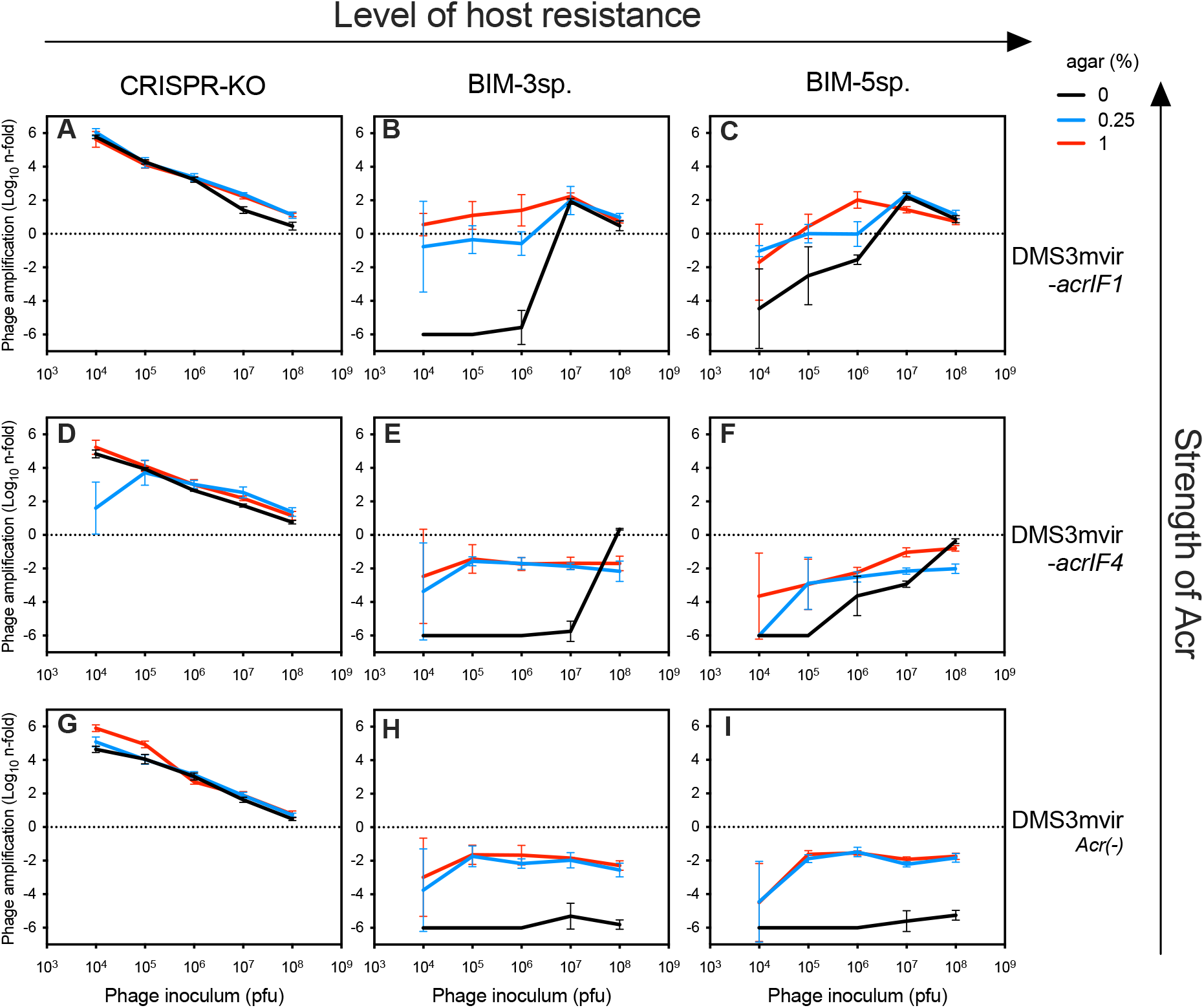
Spatial structure impacts phage epidemiology. Strains with no functional CRISPR (CRISPR-KO, A,D,G), or with 3 (BIM-3sp, B,E,H) or 5 (BIM-5sp, C,F,I) targeting spacers were infected with phage DMS3mvir without a functional Acr (Acr(-), G-I) or with a weak (*acrIF4*, D-F) or a strong Acr (*acrIF1*, A-C) introduced at different initial concentrations (corresponding to MOI ranging from 0,001 to 10). Cultures were made in M9 minimal medium with varying levels of spatial structure (from 0 to 1% agar). After one day post infection (dpi), phages were extracted and phage amplification (ratio final concentration on initial concentration) was measured. Data show the mean values for six replicates. Errors bars indicate standard deviation.

When bacteria are CRISPR-resistant, the amplification or the extinction of an Acr-phage population depends on the initial density following a phenomenon known as the Allee effect [9,17]. The phage population can either amplify if there is a sufficient number of successful infections, or reduce due to CRISPR-mediated degradation. Consistent with previous work, in well-mixed conditions, the AcrIF1-phage population amplifies only when its initial density is above 3.10^6^ PFU/ml **(Figure 1B)**. Intermediate levels of structure (blue line) do not promote phage growth, since the density threshold needed to observe an epidemic is the same as in well-mixed conditions. Note, however, that the reduction in phage concentration is less pronounced with intermediate levels of structure than in the well-mixed condition. In contrast, higher levels of structure (red line) enable the AcrIF1-phage to amplify from the smallest initial concentration tested (2,8.10^3^ PFU/ml). In other words, spatial structure can abolish the Allee effect observed in well-mixed conditions. When we carried out similar experiments with AcrIF4-phage (a phage with a weaker efficacy of Acr activity), we obtained very different results **(Figure 1E)**. In this case, spatial structure prevents the spread of the AcrIF4-phage even though the same phage can spread in well-mixed conditions.

We previously demonstrated that the Allee threshold density can be influenced by the levels of CRISPR-Cas resistance: a larger initial population of Acr-phages is necessary to invade a host population with increasing levels of CRISPR-Cas resistance [9] (compare black lines in **Figure 1B** and **1C**). We next tested if and how spatial structure affects the Allee threshold density of the AcrIF1-phage at high CRISPR-Cas resistance levels (**Figure 1C**). While increasing resistance restores an Allee effect at high spatial structure (**Figure 1C**, red line), the size of the initial phage population necessary to generate an epidemic is much smaller in structured compared to well-mixed medium (3,0.10^4^ PFU/ml versus 3,9.10^6^ PFU/ml). Therefore, our results show that high levels of spatial structure enhance the spread of the AcrIF1-phage, even when the host carries high levels of resistance. In contrast, spatial structure has no impact on the spread of the weaker AcrIF4-phage on a highly resistant bacterial host population (**Figure 1F**).

Spatial structure has thus opposite effects on two different Acr-phages. Intrigued by these ambivalent effects of spatial structure, we developed a new mathematical model to better understand the phage dynamics.

### A spatially explicit model of the dynamics of Acr phages

We developed a spatially explicit version of a model we previously used to analyse the dynamics of Acr phages in a well-mixed environment [9]. This model assumes that infection of a resistant host (R) can either lead to direct lysis and production of phage progeny (V) with a probability *ϕ* or fail and generate an immunosuppressed host (S), which can revert to a resistant state at a rate *γ* (**Figure 2**). Note that *ϕ* and *γ* are two components of the efficacy of the Acr, capturing the capacity of the protein to rapidly bind to and remain bound to CRISPR-Cas complexes, respectively. A secondary infection of S hosts automatically leads to lysis and virion production. In the spatial version of this model, the dynamics of the bacteria and the phage are taking place in two superimposed square lattices. Each site of the lattice of the host population is occupied by a bacterial cell (R or S) cell or empty (0). Similarly, each site of the lattice of the phage population is either occupied by a unit density of virus (noted with a *), or empty. Combining these different states yields 6 different types of sites (R*, S*, 0*, R, S and 0). Possible transitions among these six states are depicted in **Figure 2A**. Crucially, phage and host reproduction events are affected by spatial structure because reproduction events are conditional on the availability of hosts (for the phage) and empty spaces (for the host). We detail below how we model the influence of spatial structure. First, with probability *L* (for local reproduction) the phages adsorb to the host at the same site and, if the infection is successful, *B* newly produced phages are evenly distributed in the focal site and in the 4 neighbouring sites (**Figure 2B**). Hence, with local reproduction, phage dispersal is limited to the area around the original location of the phage. Note that, the quantity B/5 can be viewed as the unit density of phages per site and we assume there cannot be more than this quantity of phages per site. In other words, there is maximal number of phages which is set by the total number of sites on the lattice. Second, with probability 1 − *L*, the phages infect a randomly selected host in the whole lattice and, if lysis is successful, B newly produced phages are distributed randomly throughout the lattice (**Figure 2B**). Similarly, local and global host reproduction take place with probabilities *L* and 1 − *L*, respectively (**Figure 2B**, bottom panel). In other words, for simplicity we assume that the degree of spatial structure (governed by the parameter *L*) affects in the same way the dispersal kernels of the phage and its host. We analysed the individual-based model with pair approximation [18] to convert this spatial model into a set of 21 ordinary differential equations (ODEs) representing the dynamics of the different pairs of sites(detailed in Supplementary Text). We confirmed that our pair approximation captures very well the dynamical behaviour of our model with a comparison to the results of Monte Carlo simulations of the full individual-based model (**Figure S2**).

**Figure 2.**
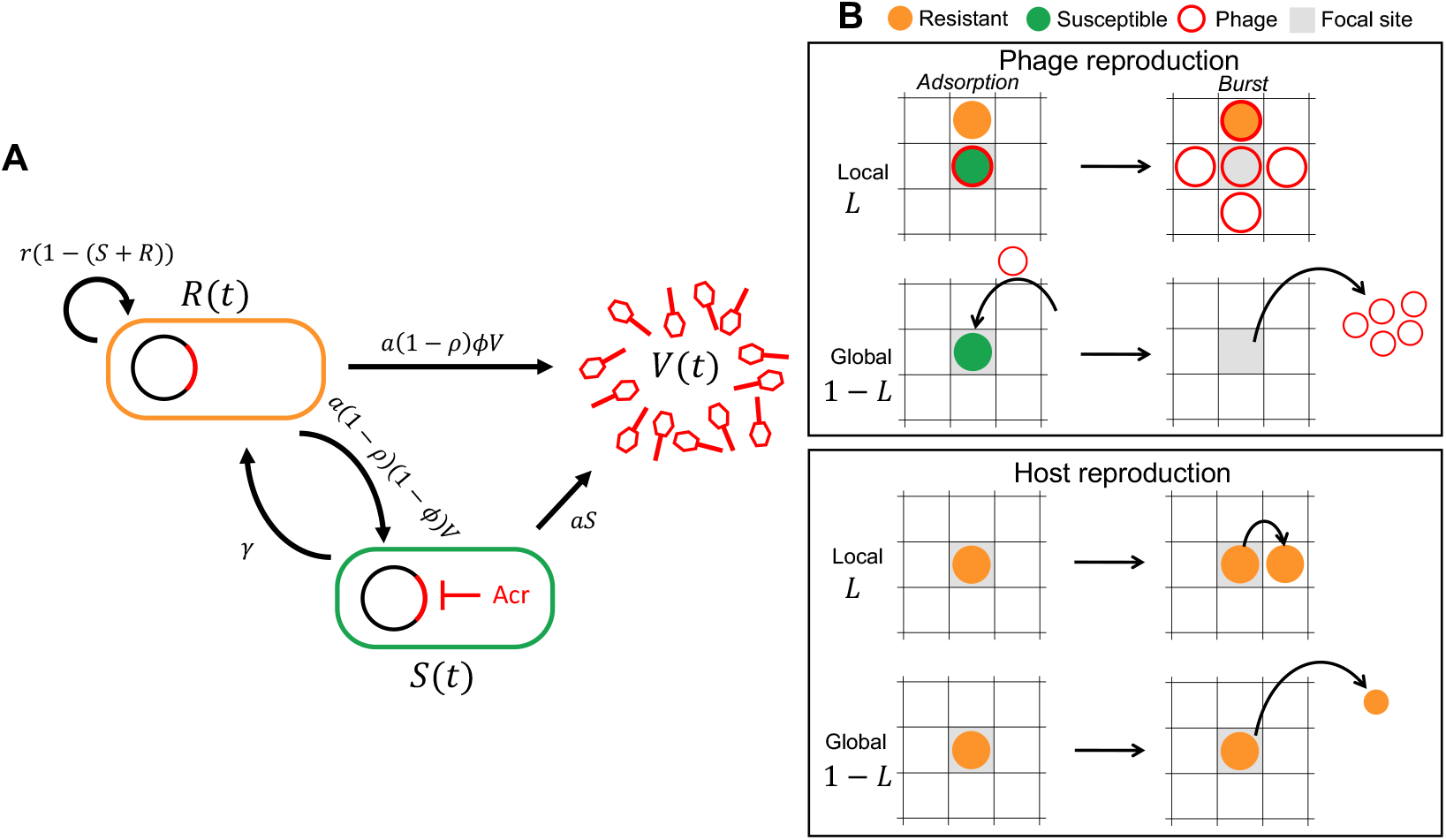
Life cycle of Acr phage and spatial dynamics. (A) A graphical diagram for the Acr-phage life-cycle showing the transition between resistant hosts (R, immunosuppressed hosts (S) and free viruses (V). Some phages manage to lyse CRISPR-resistant host cells, while the rest of the phages fail to lyse the cell and are degraded by CRISPR-resistance. Among the cells that successfully prevent lysis, a fraction of them is affected by the Acr activity of the previous phage, inhibiting CRISPR-resistance. In contrast, the infection of immunosuppressed cells always yield productive infections. (B) A graphical diagram to summarize how we model spatial structure. An example of a burst event (phage reproduction after host lysis) and a host reproduction event in our simulation is shown. In the local scenario, phages and hosts reproduce to the neighbouring sites while they disperse onto the whole lattice in the global case.

We use this model to explore the effect of the parameter *L* (a measure of spatial structure) on the phage density threshold necessary to initiate an epidemic. When *L* = 0 (well-mixed environment), phage amplification exhibits bistablity and the Acr-phage spreads only if the initial density is sufficiently high (see also [9]). In contrast, when *L* = 1 and spatial structure is high, the model predicts that Acr-phage can always invade, even from low initial densities (**Figure 3**).

**Figure 3.**
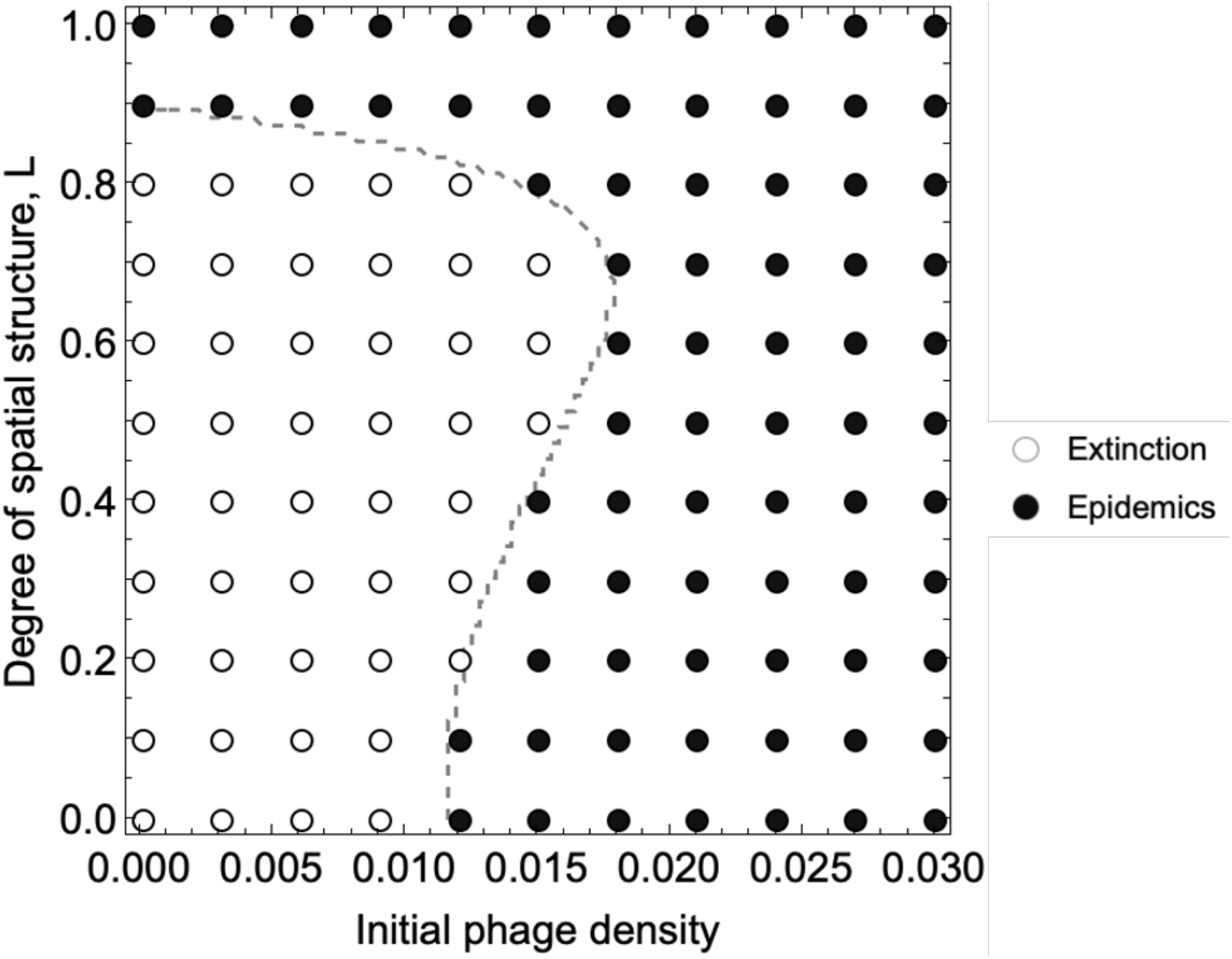
Effect of spatial structure and efficacy of Acr on epidemiological bistability. We plot the epidemiological outcome as a function of initial phage density (horizontal axis) and the amount of local reproduction *L* (vertical axis), predicted by pair approxmaion. Empty circles represent phage extinction while black dots indicate phage epidemics. The dashed line highlights the frontier between these two dynamical outcomes. This plot is based on the analysis of pair-density dynamics. Parameter values and initial values; *a* = 1, *r* = 4, *d* = 0.1, *B* = 5, *ρ* = 0.2, *γ* = 0.02, *ϕ* = 0.22, *R*(0) = 0.9, *S*(0) = 1 × 10^™14^. *V*(0) is varied from 0.01 to 0.03. The initial pair-densities are determined by the product of the initial densities of the site (R(0), S(0), V(0)).

To investigate this effect, we derive an equation for the growth dynamics of the phage (**Box 1**), which allows us to look at the early stage of the phage epidemic and to identify the effect of spatial structure on phage growth. This analysis (detailed in **Box 1** and **Supplementary Text II**) shows that spatial structure can have three effects on the growth rate of the phage population. First, it promotes phage growth by favouring the colocalization of phages and immunosuppressed hosts (*spatial exploitation effect*). Second, spatial structure also favours the spatial isolation of phages from CRISPR-resistant bacteria, which prevents them from being degraded (*spatial refuge effect*). Third, spatial structure leads to a crowding effect where new phages are more likely to land on already infected areas, limiting further spread (*self-shading effect*). The net effect of spatial structure on phage growth therefore is determined by the balance of the above three effects, which is influenced by the other parameters.

## Box 1

**Modelling the spatial spread of Acr phages**

The total phage density is the sum of all the sites occupied by a virus, 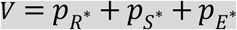, and the instantaneous growth rate *λ* of the virus results from the balance between *Adsorption* (phage loss by adsorption to *R* and *S* cells) and *Reproduction* (production of new phage after adsorption and cell lysis):

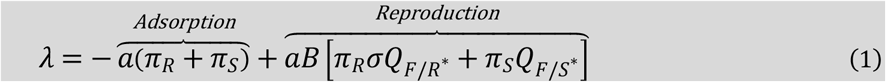

where *π*_*R*_ and *π*_*S*_ refer to the probability that the phage resides in a site occupied by a resistant bacterium (*R*) or an immunosuppressed bacterium (*S*), respectively.

- *Adsorption* to *R* bacteria may lead to three outcomes: i) with probability *ρ*, the cell is unaffected and the phage is eliminated, ii) with probability *σ* = (1 − *ρ*)*ϕ*, the host cell is lysed and *B* new phages are produced, iii) with probability 1 ™ *σ*, the phage does not lyse the cell but induces immunosuppression. *Adsorption* onto *S* bacteria always results in host lysis and the production of *B* new phages.
- *Reproduction* depends on the availability of phage-free sites in the “extended” neighbourhood of the focal site (the focal site itself and *B* ™ 1 additional sites among the nearest neighbours of the focal site). The quantities 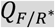 and 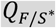 refer to the probabilities that a site in the extended neighbourhood will become a new phage-occupied site after the burst of an infected cell in the *R*\* or *S*\* site, respectively.

### Condition for the phage invasion

Equation (1) allows us to analyse the effect of various parameters on the ability of the phage to invade the host population (i.e., *λ* > 0). In the well-mixed scenario (*L* = 0), the state of the sites occupied by the phage is equal to the global state densities, (*π*_*R*_ = *R, π*_*S*_ = *S*) and the density of sites that are available is equal to the density of vacant sites 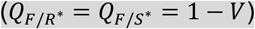. Besides, at the onset of the invasion, the phage is initially rare and all hosts are resistant (i.e., *V* ≪ 1, *R* = 1 and *S* = 0). Hence, phage invasion is only feasible when Acr efficacy is above the threshold *ϕ*) (see also [9]):

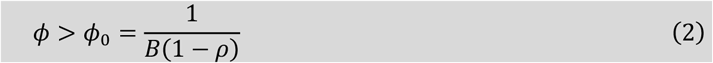

In a spatially structured population (*L* > 0), the invasion condition becomes:

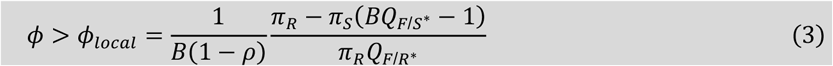

This condition is implicit because the right-hand side may also depend on *ϕ* in the local case. Yet, this new threshold provides important insights on the effects of local dispersal on phage invasion condition via 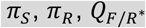 and 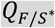. Each of these three effects have distinct consequences on the invasion success of the phage (see also **Supplementary Text II**):

#### 1 Spatial exploitation effect

Spatial structure increases *π*_*S*_, the colocalization between the phage and the immunosuppressed cells. This effect decreases *ϕ*_*local*_ and promotes phage invasion because it increases access to susceptible cells,

#### 2 Spatial refuge effect

Spatial structure reduces *π*_*R*_, the colocalization between the phage and the resistant cells. This effect decreases *ϕ*_*local*_ and promotes phage invasion because it reduces the contact with resistant cells.

#### 3 Self-shading effect

Spatial structure decreases 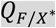 via phage clustering. This effect increases *ϕ*_*local*_ and hinders phage invasion because it limits diffusion to phage-unoccupied sites.

These antagonistic effects capture the ambivalent effect of spatial structure and our numerical simulations show that the balance between these opposite forces is driven by the persistence of immunosuppression. Spatial structure promotes phage invasion (*ϕ*_*local*_ < *ϕ*_0_) when immunosuppression is long. But spatial structure has the opposite effect (*ϕ*_0_ < *ϕ*_*local*_) when immunosuppression is short (Figure 4).

**Figure 4.**
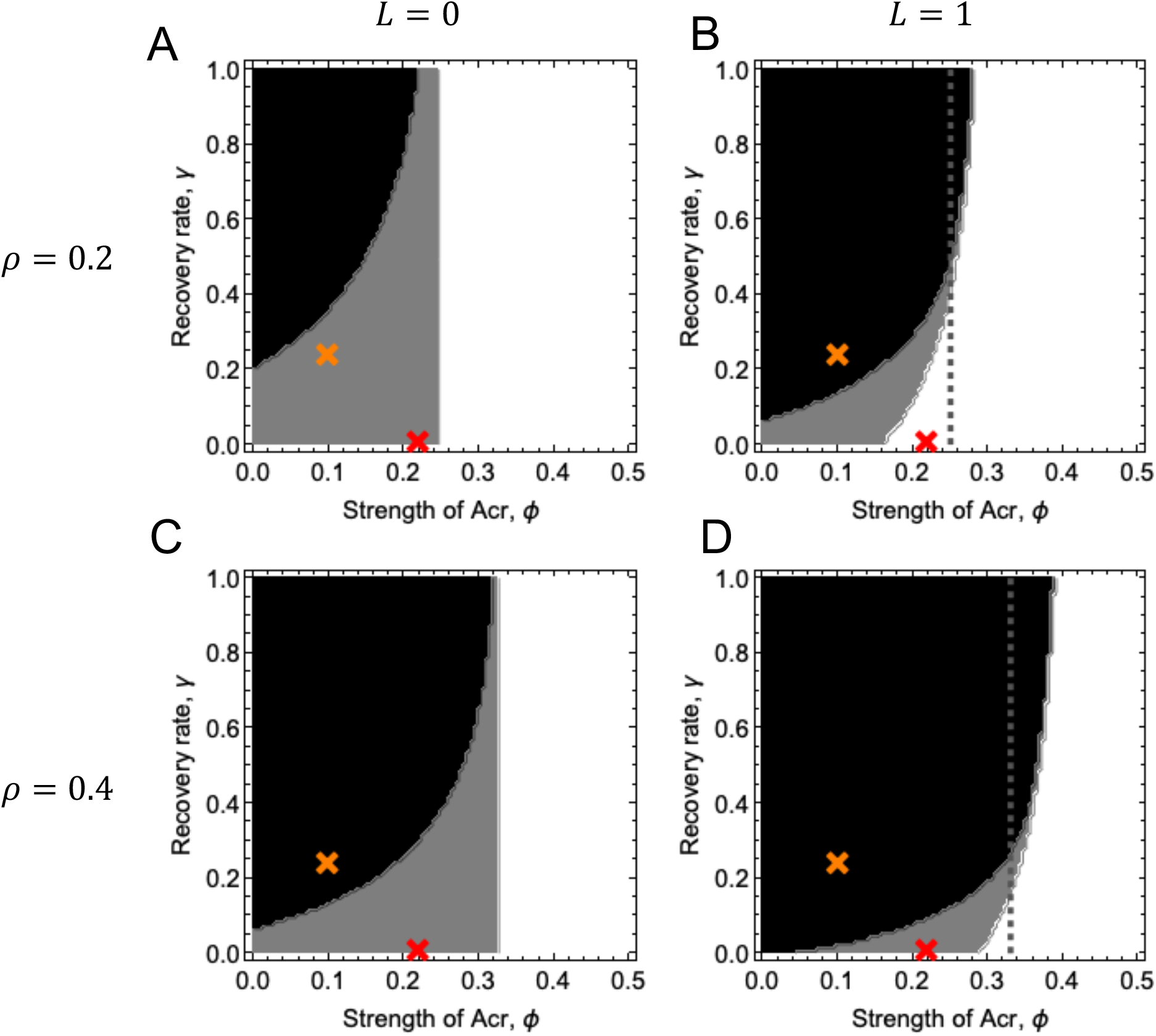
Epidemiological outcomes for well-mixed and structured populations. Epidemiological dynamics based on pair-density dynamics are analysed on the combination of Acr efficacy (*ϕ* and *γ*). Left panels (A,C) shows the outcomes in the well-mixed population while the right panels (B, D) show those in highly structured populations. In upper panels (A, B), phages challenge the weak CRISPR-host population (*ρ* = 0.2) while they challenge the stronger hosts (*ρ* = 0.2) in the lower panels (C, D). Black area shows phages do not cause epidemics while white area phages always cause epidemic. Gray area shows the epidemiological bistability where the outcome depends on the initial conditions. The vertical line in (A, C) and the dashed vertical line in (B,D) is *ϕ*_)_ in Box 1. The curve that separates grey and white area in (B, D) is *ϕ*_*local*_ in Box1. Coloured crosses refer to phages with different efficacy of Acr (red: (*ϕ, γ*) = (0.22,0.02), orange: (*ϕ, γ*) = (0.1,0.25)). Parameter values and initial values: *a* = 1, *r* = 4, *d* = 0.1, *B* = 5, *ρ* = 0.2, *γ* = 0.02, *ϕ* = 0.22, *R*(0) = 0.9, *S*(0) = 1 × 10^™14^, *V*(0) = 0.0001 The initial pair-densities are determined by the product of the initial densities of the site (R(0), S(0), V(0)).

To understand the effect of spatial structure on the initial dynamics of Acr-phage, we contrast two extreme scenarios in **Figure 4** with no spatial structure (*L* = 0), or with a maximal level of spatial structure (*L* = 1). In each scenario, we vary the efficacy of CRISPR-resistance (the parameter *ρ*) and the efficacy of Acr (the parameters *ϕ* and *γ*) and three different outcomes are possible: (i) the phage can spread when initially rare (white area in **Figure 4**), (ii) the phage cannot spread when initially rare, but can spread when the initial density of the virus is high (grey area), (iii) the phage can never spread (black area).

## The spread of Acr-phage depends on the efficacy of CRISPR-resistance and Acr activity

Our theoretical analysis shows that higher CRISPR-resistance (*ρ*) restricts phage growth and narrows the conditions under which phages can successfully invade, in both well-mixed (**Figure 4A** vs. **4C**) and spatially structured environments (**Figure 4B** vs. **4D**). On the other hand, a higher probability of direct lysis by Acr-phage (*ϕ*) consistently enhances phage growth across all conditions. Notably, in well-mixed settings, the invasion threshold for *ϕ* is independent of the duration of immunosuppression (*γ*), whereas under spatial structure, this threshold increases with *γ*. Furthermore, longer immunosuppression (i.e., smaller *γ*) always favours phage growth by extending access to susceptible host cells.

These theoretical insights are consistent with experimental observations. Indeed, increased levels of CRISPR-resistance reduce Acr-phage amplification in both well-mixed and structured environments, as shown by fewer data points above the no-amplification line in **Figure 1C, F**, and **I** compared to **Figure 1B, E, and H**. Moreover, the strong AcrIF1-phage consistently outperforms the weaker AcrIF4-phage in initiating epidemics under all tested conditions (**Figure 1A–C** vs. **1D–F**), confirming the critical role of Acr strength and host resistance in determining invasion success.

Importantly, our theoretical analysis allows us to explore the effect of spatial structure and to better understand how the efficacy of Acr interacts with this effect and may generate the ambivalent effect we observed experimentally.

## The effect of spatial structure on the bistability of Acr-phages depends on the efficacy of Acr

The bistability (grey area) is reduced with spatial structure (compare **Figures 4A** with **4B** and **4C** with **4D**; see also **Figure S5**). The effect of the initial density of the virus in well-mixed populations is only driven by its influence on the initial density of immunosuppressed hosts [9]. In spatially structured populations, the initial density of the virus interferes with the effect of local dispersal on invasion condition via additional effects (**Box 1**). This effect of spatial structure on phage invasion depends also on the efficacy of Acr. When Acr efficacy is relatively weak (orange cross in **Figure 4**), spatial structure reduces invasibility of phages (change from grey to black between **Figures 4A** and **4B**). In contrast, when Acr efficacy is relatively strong (red cross in **Figure 4**), spatial structure allows the phages to invade even when rare (change from grey to white between **Figures 4A** and **4B**).

The above results may thus help understand the ambivalent effect of spatial structure we observed experimentally. **Figure 1** shows that increasing spatial structure can reduce Acr-phage bistability when Acr is strong and also reduces it when Acr is weak. Indeed, high levels of spatial structure abolished the Allee effect during the infection of BIM-3sp bacteria (intermediate resistance) by the strong AcrIF1-phage (**Figure 1B**, red line always above the dotted line), and therefore strongly favoured AcrIF1-phage growth. In contrast, we noticed that spatial structure disfavoured the spread of the weak AcrIF4, as phage amplification on BIM-3sp bacteria could only occur in well-mixed conditions, but not in structured medium (**Figure 1E**).

## The effect of spatial structure on the spread of Acr-phages depends on the duration of immunosuppression

When immunosuppression is short-lived (large values of *γ*) spatial structure reduces the growth rate of the phage. In contrast, when immunosuppression can persist (low values of *γ*), spatial structure increases the ability of the phage to invade when rare (even though bistability might be reduced). To understand the interaction between spatial structure and immunosuppression we can examine the threshold value *ϕ*_*local*_ derived in **Box 1**. At the beginning of the invasion of the Acr-phage, the threshold *ϕ*_*Local*_ is above *ϕ*_*Global*_,which indicates that spatial structure is hindering the spread of the phage (**Figure S4**). Indeed, in a spatially structured environment, the phage has only access to a fraction of the host population, and this limits the initial growth rate of the phage (**Supplementary text II-; Figure S3**). Yet, this effect is only transitory and *ϕ*_*Local*_ reaches a maximum around *t* = 2 and drops afterwards. Crucially, the magnitude of the drop of *ϕ*_*Local*_ depends on the persistence of immunosuppression. When immunosuppression is short-lived (large values of *γ*) this drop is reduced and *ϕ*_*Local*_ remain higher than *ϕ*_*Global*_. In contrast, when immunosuppression is long-lived, *ϕ*_*Local*_ drops below the threshold value *ϕ*_*Global*_. In these situations, spatial structure increases the local density of immunosuppressed hosts in the neighbourhood of the phage (**Supplementary text II-; Figure S3**) which boosts its growth rate.

Interestingly, the main parameter driving the difference between AcrIF1 and AcrIF4 is thought to be the duration of immunosuppression [16] (see **Supplementary text III**). Hence, the short-lived immunosuppression (high *γ*) of AcrIF4 is consistent with its inability to generate an epidemic when spatial structure becomes high (orange cross transitions from grey to black area in **Figure 4A, B** may explain **Figure1E**). In contrast, the long-lived immunosuppression (low *γ*) of AcrIF1 enable their unconditional amplifications in highly spatially structured population (see red cross, which changes from grey to white area in **Figure 4C, D**, may explain **Figure 1B**).

## Discussion

Phage-bacteria interactions are commonly studied in liquid and well-mixed cultures where their encounter rate are proportional to the densities of their populations. In such a homogenous population each cell experiences the same risk of infection, which is reflected by the global multiplicity of infection (MOI). In contrast, most natural environments are spatially structured and different cells can be exposed to very different risks of infection. In particular, the MOI can vary in space and can be locally very high. This heterogeneity can have strong influence on phage epidemics and bacterial survival. Typically, spatially structured environments were often found to protect bacteria by slowing down the spread of phage epidemic [19,20]. Indeed, phage proliferation is limited to a local area while bacteria further away from this area are inaccessible to phages. These spatial refuges can be generated by the spatial organisation of the gut epithelium or by bacterial growth within a biofilm or as a colony [21–27]. Although spatially structured environments limit phage proliferation, they were found to generally promote the coexistence of phages and susceptible bacteria [21–28].

In contrast to this spatial refuge model, the present study examined whether spatial structure could enhance the propagation of Acr-phages that rely on cooperative behaviours to multiply [9,10] and found compelling evidence that it does. Notably, we assessed how the epidemiology of Acr-phages is influenced by the interactions between spatial structure and three parameters that were previously showed to modulate Acr-phage amplification, namely i) the initial phage density, ii) the level of CRISPR-resistance and iii) the inhibitory efficacy of the Acr protein [9]. Our experiments indicate that spatial structure can abolish the characteristic bistable growth of phage populations that rely on group behaviour (Allee effect) and promotes phage propagation from low population densities. However, this positive effect of spatial structure disappears when the level of host resistance to phages is increased or when the capacity of the phage to inhibit resistance is less efficient. In this latter case, the effect of spatial structure on phage fitness is even reversed: phages that normally grow in well-mixed conditions no longer propagate in a structured environment.

This ambivalent effect of spatial structure was unexpected, because spatial structure increases locally the MOI and should promote the spread of the Acr-phages. To better understand the causes of this ambivalence, we developed a spatially explicit mathematical model. Our model brings new insights on the effect of spatial structure and extends the theoretical framework of spatial evolutionary epidemiology [20,29–31]. The ambivalent effect of space results from the presence of different types of hosts with different productivities in the population (S and R hosts). Hence, spatial structure acts on the encounter rates between the virus and the two types of hosts (S and R) as well on the intensity of competition among viruses. Spatial structure exerts three key effects on phage dynamics (see **Supplementary Information**): it enhances access to susceptible hosts (spatial exploitation effect), reduces contact with resistant hosts (spatial refuge effect), and limits effective reproduction by redirecting phages to already infected sites (self-shading effect). The overall influence of spatial structure on phage dynamics results from the sum of these three effects. In particular, we found that the duration of Acr-mediated immunosuppression is key to understand why spatial structure has an ambivalent effect on the growth of Acr-phages: Acr that generate short-lived immunosuppression (weak Acr) are disfavoured while Acr that induce long-lived immunosuppression (strong Acr) are strongly favoured. Indeed, when immunosuppression lasts longer, it becomes easier for susceptible cells to reach higher densities. Our findings align well with theoretical predictions concerning the interplay between spatial structure and population dynamics under Allee effects. In particular, the work of Surendran et al. (2020) highlights how local interactions can fundamentally alter extinction thresholds in populations exhibiting strong Allee effects [32].

Spatial structure is often viewed as a key driver in the evolution of cooperation. Can the ambivalent effect of space on viral spread affect the evolution of cooperation in the virus population? Our previous work in well-mixed conditions suggested that Acr with short duration of immunosuppression provides a larger competitive advantage than Acr proteins with long immunosuppression time during competition with phages lacking Acr. This is because strong Acr-phage share their benefits with phages lacking Acr, helping these “cheater” phages to reproduce on CRISPR-resistant bacteria, while weak Acr-phage do not and therefore have a more “selfish” behaviour [16]. Hence, weak Acr-phages may be better adapted to well-mixed environments, where the benefits of Acr production are confined to the producing phages themselves, thereby minimizing the risk of exploitation by cheaters. In contrast, strong Acr-phages, which are more vulnerable to exploitation, may thrive in spatially structured settings that may offer protection against cheaters. Therefore, spatial structure may favour the evolution of Acr proteins conferring strong inhibition of CRISPR-Cas system. A recent theoretical work by Segredo-Otero and Sanjuán showed that spatial structure can promote the evolution of cooperative immune evasion strategies in viruses [33]. In their model, inhibition of interferon secretion - a cooperative trait [34] - can be favoured by spatial structure because it enhances the clustering of cooperators and limits exploitation by cheaters. These insights support the idea that spatial structure plays a critical role in the evolution of cooperative behaviours among viruses. Future investigations into whether strong Acr-phages remain susceptible to exploitation in spatially structured environments will clarify whether similar dynamics arise and provide valuable insights into the evolutionary implications of spatial structure. Note, however, that since our experimental model did not reveal any detectable fitness costs associated with Acr production, it remains unclear whether cheating could indeed disrupt cooperative interactions among Acr-phages [16]. More generally, the importance of our results for other Acr-phages remain to be investigated given that only few Acr-phages were found to rely on cooperation. Besides, it would be interesting to explore if, beyond the adaptation of phage to CRISPR-resistance, phage can cooperate to escape other host defence systems. Yet, our joint experimental and theoretical study clarifies the interplay between spatial structure, population dynamics and cooperative behaviour on the spread of a virus population. This analysis confirms that spatial structure is a key environmental factor that needs to be accounted for to predict the epidemiology and evolution of pathogens.

## Methods

### Experimental procedures

#### Bacteria

Throughout this study we used strains derived from the wild-type strain UCBPP-PA14 of Pseudomonas aeruginosa. Strains BIM-3sp and BIM-5sp carry 3 or 5 spacers targeting phage DMS3mvir, respectively and the strain CRISPR-KO, (UCBPP-PA14 *csy3::lacZ*) has a non-functional CRISPR-Cas system. Detailed descriptions of these strains can be found in Landsberger et al. (2018) and references therein. Bacteria were cultured at 37°C in M9 minimal medium (22 mM Na_2_HPO_4_; 22 mM KH_2_PO_4_; 8.6 mM NaCl; 20 mM NH_4_Cl; 1 mMMgSO_4_; 0.1 mM CaCl_2_) supplemented with 0.2% glucose (referred to as M9 medium in the rest of this manuscript).

#### Phages

The Mu-like virulent phage DMS3mvir and isogenic variants carrying anti-CRISPR genes, namely DMS3vir-*acrIF1* and DMS3vir-*acrIF4*, were used throughout this work (described in [9,35,36]). Phage stocks were obtained from lysates prepared on PA14 CRISPR-KO and stored at 4°C.

#### Infection Assays

Vials containing 3 mL of M9 medium, M9 0.25% agar or M9 1% agar, were prepared the day before the experiment and kept at 60°C. These pre-filled 15 mL-falcon tubes were placed at 42°C 5min prior inoculation to allow temperature equilibration while avoiding that the M9 agar 1% medium sets. Each vial was rapidly inoculated with approximately 10^7^ colony forming units (CFU) from fresh overnight cultures of CRISPR-KO or BIM-3sp or BIM-5sp strains and with 10^8^ or 10^7^ or 10^6^ or 10^5^ or 10^4^ plaque forming unit (PFU) of phage DMS3mvir or DMS3mvir-*acrIF1* or DMS3vir-*acrIF4*. The vials were thoroughly mixed straight away after inoculation (vortex) and incubated at 37°C under agitation. One hundred microliters were sampled from vials with M9 medium, serially diluted and plated on LB agar and spotted on lawns of the CRISPR-KO strain to verify the bacterial and phage titres, respectively.

Upon 24h-incubation, 3mL of warmed M9 medium (42°C) were added to each vial to help agar disruption and homogenise the medium (vortexing). For vials with 1% agar, further treatment consisting in mechanical disruption and vigorous shaking were necessary to be able to pipet. We then added 10% chloroform, vortex for 15 s and centrifuge (20 min, 3500 rpm, 4°C) all vials. Samples were collected, serially diluted in buffer and spotted on lawns of CRISPR-KO to determine the phage titre 1 day post infection.

In summary, 24h-infection assays were performed while varying 4 parameters (3 agar concentrations, 3 bacterial strains, 3 phage strains, 5 phage inoculum) and each assay was performed in 6 replicates. A total of 810 infection assays were performed.

#### Theory

In this section, we describe a dynamical model of host bacteria with CRISPR-resistance system (phage-resistant bacteria) and Acr-phages (anti-resistant phages) that have the countermeasure against CRISPR-resistance. Our focus is on the effect of spatial structure on the epidemiological dynamics of phage-resistant bacteria and anti-resistant phages. We first introduce the model in the absence of spatial structure and then incorporate spatial dynamics using a dual lattice structure with two overlayed lattice (one for the bacteria population and one for the phage). In addition to individual based simulation, we use pair approximation [18,37–39] to analyse the epidemiological dynamics of phage and bacteria in spatially structured populations.

#### Well-mixed model

Before introducing the spatial structure, we begin with the epidemiological dynamics of Acr phage in a well-mixed bacterial population, as proposed in a previous study [9]. Let us consider the situation where Acr-phage is introduced into the host population where all hosts are initially resistant to phage infection due to CRISPR-resistance (i.e. all bacteria already have spacer sequence for the phage). The epidemiological dynamics of Acr-phage in this host population is described as follows.

#### Phage infection

After the phage adsorbs on the resistant bacterial cells, three different outcomes may occur. (1) The phage may fail to infect with probability *ρ* due to CRISPR-resistance. If infection is successful, which occurs with probability 1 − *ρ*, either (2) phage lysis occurs with probability *ϕ*, or (3) lysis does not occur with probability 1 − *ϕ*. In the latter case, the phage-encoded Acr protein inhibits CRISPR-resistance of hosts, thereby making them temporally susceptible to the phage. Susceptible hosts revert to immunized hosts at a rate *γ* through inactivation of Acr. When the phage adsorbs on the susceptible bacterial cell, in contrast, the cell always lyses.

#### Phage reproduction

A single phage lysis gives rise to *B* newly produced phages. Although not assumed in the previous model [9], we here assume that there is a carrying capacity for phages. Note however that this additional assumption does not change the invasibility of Acr-phage, which is the main topic of the present, as well as previous paper [9], because phage density is sufficiently small in the early stages of the epidemic.

The infection process described above is combined with the process for bacterial births and deaths to yield the following time changes in the densities *R* and *S* of resistant (CRIPPR immunized) and susceptible (immunosuppressed) host bacteria and the density *V* of Acr phages:

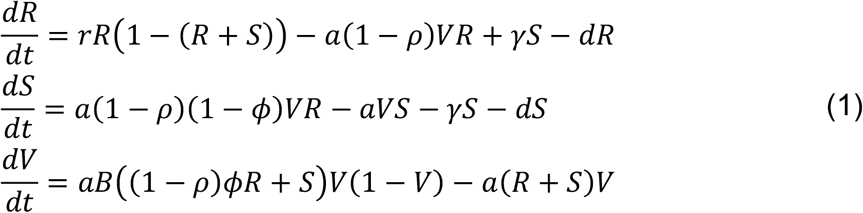

where *r* is the reproduction rate, *d* is the death rate of host bacteria, *a* is the adsorption rate and *B* is the burst size of phages. In this study, the (scaled) burst size *B* is set to 5 to make the comparison with the spatially explicit (dual-lattice) model easier, where, in the latter, newly produced phases after lysis are sent to the focal site as well as 4 nearest neighbor sites (*B* = 1 + 4). *γ* is the rate of transition from *S* to *R* (loss of immunosuppression). The coefficient of density-dependent mortality of bacteria is scaled to 1. It is assumed that immunosuppressed bacteria do not reproduce.

#### Spatially explicit model

The above well-mixed model is extended to a spatially explicit model in which both bacteria and phage inhabit sites on a square lattice. Bacteria send its progeny to the empty (bacteria-free) site in the nearest neighbours. After lysis, newly produced phages are sent to the focal and its 4 nearest neighbour sites if they are phage-free. Phage adsorption and reversion to resistance, as well as host natural death, occur spontaneously within the focal site (**Fig. 2**). Each site of the lattice can be occupied by at most one host and one unit density of phages. In the host lattice, each site is occupied by *R, S*, or empty. In the phage lattice, each site is either occupied by one unit density of phages or empty. Therefore, there are six different states of each site denoted by *R*\*, *S*\*, 0*, *R, S*, and 0,where *R* and *S*denotes respectively the site which is phage-free but occupied by resistant and sensitive bacteria, while 0 denotes the site which is both phage-free and bacteria-free. The sites *R*\*, *S*\*, and 0* denotes respectively the corresponding sites but with phage particles (**Fig. 2B**). We consider that the transition among these six states follows the life cycle detailed above. Phage adsorption occurs at sites where both the phage and the host are present (i.e. *R*\* and *S*\*). Among the events that change the state of a site, both the phage and the host reproduction are particularly affected by the spatial structure (**Fig. 2B**). We assume that both phage and host reproduction occur either *locally* with probability *L* (i.e., phages are adsorpted to the hosts in the same location and newly produced phages and newly produced hosts are sent to the nearest neighbouring sites) or *globally* with probability 1 − *L* (i.e., phages are adsorpted to any sites and the newly produced phages and hosts are sent to a site randomly picked in the whole lattice).

If the new site is already occupied by a virus, the new site keeps its previous state (i.e. infected). Similarly, if the new site is already occupied by a host, the new site keeps its previous state. In other words, we assume there can be at most one phage and one host per site.

#### Pair approximation

To analyse the spatial epidemiological model presented above, we used pair approximation to account for spatial correlations over space by local pair densities [18,37]. Let *p*_*x*_ be the proportion of sites in state *x* (*x* ∈ *G* = {*R*\*, *S*\*, 0*, *R, S*, 0}). Let *p*_*xy*_ be the doublet density of a nearest neighbor pair *xy* in the population, and *q*_*x*/*y*_ = *p*_*xy*_/*p*_*y*_ be the local density of *X* site in the local neighborhood (i.e., the 4 nearest sites) of sites in state *Y* (*x, y* ∈ *G*). Also, let *q*_*x*/*yz*_ = *p*_*xyz*_/*p*_*yz*_ be the conditional probability that a site chosen randomly in the local neighborhood of *y* in the *yz* pair is in state *x* site, where *p*_*xyz*_ is the density of triplet *xyz*. Pair approximation is a method to decouple tertial or higher spatial correlations of spatially explicit models by assuming that *q*_*x*/*yz*_ = *q*_*x*/*y*_.

As explained in the **Supplementary Information**, the well-mixed population model (1) can be recovered from the pair density dynamics by applying mean field approximation that assumes *q*_*X*/A_ = *p*_*X*_, where *p*_*X*_ is the density of singlet *X*, and denoting 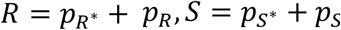, and 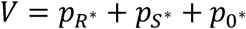.

As noted above, spatial structure has a significant impact on Acr phage epidemics. Pair approximation can capture the effect of spatial structure on the phage dynamics shown in Monte Carlo simulations of the dual lattice model (see **Supplementary Information** for details), and provide significant insights into the experimental results for the spread of Acr-phages in CRISPR-resistant bacteria.

